# Limitations of general protein language models and public TCRpMHC data in training specificity prediction models

**DOI:** 10.64898/2026.09.21.753204

**Authors:** Sarah Hall-Swan, Brajesh Rai

## Abstract

T cell receptor (TCR) specificity prediction is critical for understanding adaptive immunity and for enabling the development of therapeutic interventions. Current machine learning models for predicting TCR-peptide-MHC interactions perform poorly, despite large efforts by many groups and advances in protein language models. This work identifies and addresses the fundamental issues that might be hindering progress in this field, focusing on current approaches for constructing training and validation sets, particularly on the strategies used by commonly used methods, which generate negative data either by shuffling TCRs between epitopes or by pairing epitopes with TCRs from background datasets of healthy donors. Here, we visualize and quantify the overlap between positive and generated negative TCR sets produced by both strategies using a range of sequence embedding approaches, including simple encodings (one-hot and BLOSUM62), general protein language models (ESM-2 and ProtGPT2), and a TCR-specific embedding model (TCR-BERT). Using paired *αβ*TCR data from VDJdb for two well-studied epitopes (GILGFVFTL and KLGGALQAK), we train Random Forest classifiers on each embedding representation and evaluate their ability to distinguish positive from negative data examples. We find that general sequence embedding methods struggle to differentiate between positive and negative TCRs. We further show that the choice of negative data generation method significantly impacts classifier performance. Our findings highlight fundamental limitations in current TCR specificity training data and general protein language models, and underscore the need for TCR-specific embeddings, experimentally validated negative datasets, transparent reporting of negative data construction, and epitope-specific modeling approaches.

## Introduction

The adaptive immune response provides a primary defense against pathogens. It is composed of humoral immunity, mediated by B cells, and cellular immunity, mediated by T cells. In cellular immunity, peptides generated within cells are presented on the cell surface by major histocompatibility complex (MHC) molecules as peptide-MHC (pMHC) complexes. T cells recognize foreign or mutated peptides bound to MHCs through specialized T cell receptors (TCRs), enabling cytotoxic T cells to eliminate infected cells and helper T cells to coordinate and amplify immune responses^1^. Accurate prediction and characterization of TCR-pMHC interactions are therefore central to understanding adaptive immunity and has important therapeutic applications. Vaccines can be developed to trigger helper T cells and create an immune response that will activate when the individual is infected by a specific virus^2,3^. Certain kinds of cancer immunotherapy, such as CAR T cell therapy, are developed to enhance the adaptive immune response against cancer cells^4^. Autoimmune disease can be the result of T cells mistakenly targeting self-peptides, thus harming healthy cells. Therefore, predicting TCR specificity can aid in preventing autoimmune reactions^5^.

TCR specificity prediction is made complicated by the high diversity of the TCR repertoire. An individual human is estimated to possess approximately 2x 10^6^ distinct TCRs in the naive TCR repertoire^6^. This diversity arises from somatic recombination of the variable (V), diversity (D), and joining (J) gene segments that make up the variable domain of the TCR*β* chain, and the V and J genes of the *α* chain. The *α* and *β* chains subsequently pair to form the *αβ*TCR (the same process is carried out for the less common *γδ*TCR). TCR diversity is necessary for effective immune defense, both in quickly responding to a newly introduced peptide and allowing for immune flexibility to guard against pathogen mutations^7^. However, when combined with the high number of potential peptide-MHC complexes, this diversity makes comprehensive experimental characterization of all possible TCR-pMHC interactions infeasible. Therefore, machine learning approaches are increasingly used to predict TCR specificity based on patterns in existing datasets. The capabilities of these machine learning methods critically depend on the size and diversity of available training data.

Several public repositories storing TCR specificity data are available. The “completeness” of the available data ranges from containing only CDR3*β* regions, to sequences with full paired *α* and *β* chains of the TCR. One such repository is VDJdb^8^, which curates TCR-pMHC binding data from published studies and includes both paired and unpaired TCR chain information. The vast majority of epitopes in VDJdb are associated with class I MHC molecules, with a smaller number bound to class II MHCs. Another manually curated data repository, McPAS-TCR^9^, similarly contains paired and unpaired TCR chain data. In both VDJdb and McPAS-TCR, entries may only contain CDR3 regions or include the V and J genes as well. The ImmuneCODE database contains over 160,000 SARS-CoV-2 antigen-associated TCRs identified using the MIRA protocol. However, this dataset is limited to TCR *β* chains^10^. Finally, the Immune Epitope Database (IEDB) includes results from MHC, B cell, and T cell assays^11^, and unlike other repositories, contains some experimentally derived negative T cell assay entries, although these often lack corresponding T cell sequence information.

Given the scarcity of experimentally validated negative TCR specificity data, most supervised TCR specificity models rely on synthetically generated negative examples. One common strategy generates negatives by shuffling TCRs between epitopes, creating negative pairs by associating TCRs with peptides to which they are not reported to bind in the positive dataset. This approach is often based on the assumption that sufficiently dissimilar epitopes are unlikely to be recognized by the same TCR^12,13^. An alternative strategy pairs epitopes from the positive dataset with TCRs drawn from a “background” repertoire from independent studies^14–19^. In both cases, the resulting negative examples are not experimentally validated as non-binding and therefore may contain false negatives.

Many TCR specificity prediction methods use the sequence of the TCR (and, in some cases, the peptide and MHC), creating a need for sequence representations that accurately capture features relevant to antigen recognition, typically in the form of a sequence embedding^14,15,17,19^. Traditional sequence representations rely on predefined embeddings, such as one-hot encoding of amino acids or substitution-matrix-based encodings derived from BLOSUM^20^ matrices. More recently, general protein language models have emerged as deep learning approaches for generating protein sequence embeddings, motivated by the success of large-language models^21,22^. Embeddings produced by these models have been adopted as inputs to TCR specificity prediction models, including TCR-ESM^23^ and STAG-LLM^16^.

In this study, we use protein language model embeddings to assess the overlap between positive TCRs and generated negative TCR sets. We find substantial overlap in the embedding space, suggesting that commonly used negative data generation strategies may be insufficient for training supervised machine learning models for TCR specificity prediction. We further compare the performance of Random Forest^24^, Gradient Boosting, Multilayer Perceptron, and Logistics Regression classifiers trained on a range of embedding representations, including simple sequence encodings (one-hot and BLOSUM62), general protein language models, and TCR-specific embeddings.

## Methods

### Positive and negative data

Positive training examples were obtained from TCR-pMHC binding assays in VDJdb^8^. We limited the data to human TCRs and MHCs, requiring paired *αβ* TCR chains and peptides presented by class I MHC molecules. We refer to this collection as the Full dataset. From this dataset, we defined two epitope-specific positive sets: one comprising TCRs that bind the influenza A-derived epitope GILGFVFTL presented by HLA-A*02:01 (the GIL set), and the other consisting of TCRs that bind the human herpesvirus 5-derived epitope KLGGALQAK presented by HLA-A*03:01 (the KLG set). The GIL and KLG sets contain 1,892 and 13,366 TCR sequences, respectively.

For each positive set, we constructed corresponding “negative” sets using two commonly employed strategies. The first approach generates negatives by shuffling TCRs across epitopes, as used in prior studies^12,25,26^. Under this scheme, TCRs in the Full dataset that bind pMHCs other than the epitope of interest are treated as candidate negatives. Accordingly, the negative set for the GIL dataset consists of TCRs in the Full dataset not reported to bind to GILGFVFTL-A*02:01, while the negative set for the KLG dataset comprises TCRs not reported to bind KLGGALQAK-A*03:01. It is important to emphasize that these negative TCRs have not necessarily been experimentally validated as non-binding; rather they are assumed to be negative based solely on their absence from the corresponding the positive set.

The second negative set was constructed for both positive datasets. We use as set of background TCRs, as done in previous studies^15,17,18^, sequenced from healthy donors, obtained from the Observed T cell Space (OTS), a curated repository of paired *αβ* TCR sequences aggregated from 50 studies^27^. The full healthy background set contains 128,340 TCRs. To avoid overlap with the positive data, any TCRs present in the positive datasets were removed from the background set.

### Full sequence extraction

From VDJdb and the OTS, TCRs are provided as V and J genes, and CDR3 amino acid sequences for the *α* and *β* chains. We used the python API of the stitchr software^28^ to construct the full amino acid sequence of the each chain, after which the a and *β* chain sequences were concatenated to generate the full TCR sequence.

### Sequence embedding

Sequence embeddings of each full TCR sequence were generated using the Evolutionary Scale Model (ESM-2)^21^ developed by the Meta Fundamental AI Research Protein Team. ESM-2 is a general protein language model trained on sequences drawn from approximately 45 million UniRef50 clusters^29^. In this study, we used the pretrained model `esm2_t36_3B_UR50.` Per-residue embeddings were extracted from the final layer of the transformer. Per-sequence embeddings are generated by averaging the per-residue embeddings after removing the padding tokens. Sequence embeddings were generated in the same manner using ProtGPT2^22^, another foundation protein language model trained on approximated 50 million sequences from UniRef50 version 2021_4^29^. As of February 2026, UniRef50 contains roughly 60 million clusters in UniRef50, fewer than 1,000 of which are annotated as T cell receptors.

Sequence embeddings were also generated using TCR-BERT, a TCR-specific embedding model trained on the CDR3 regions of TCR alpha and beta sequences from the pan immune repertoire database (PIRD) and VDJdb, comprising approximately 85,000 *β* sequences and 30,000 *α* sequences^30^. Because TCR-BERT is restricted to the CDR3 region, we used embeddings derived from the CDR3*β* region. As with ESM-2, per-sequence embeddings were obtained by averaging the per-residue embeddings after removal of padding tokens.

Lastly, to compare protein language models with simpler sequence representations, we generated embeddings using one-hot encoding of amino acid sequences and an embedding based on the BLOSUM62 substitution matrix^20^. Sequences were padded with the character “X” to a fixed length of 601. For one-hot encoding, each amino acid was represented by a 21-dimensional binary vector corresponding to its position in the alphabet “ACDEFGHIKLMNPQRSTVWYX”, and full TCR sequences were encoded by concatenating the vectors for each amino acid in the sequence. For the BLOSUM62 embedding, each amino acid is represented by a 25-dimensional vector corresponding to its row in the BLOSUM62 matrix, with the padding character “X” encoded as a vector of zeros. Full TCR sequence were similarly represented by concatenating residue-level BLOSUM62 vectors.

### Classification

To quantify the overlap between positive and negative TCRs in embedding space, we trained various classifiers, including Random Forest, Gradient Boosting, Multilayer Perceptron (MLP), and Logistic Regression to distinguish positive from negative TCRs using the sequence embeddings as input features. For each positive set (GIL and KLG), two of each classifier were trained, one for each negative set (shuffled negatives and healthy background negatives). Model hyperparameters were tuned using 5-fold cross-validation and final model performance was evaluated on a held-out test set. We report the area under the precision-recall curve (AUPRC) for each classifier.

We used the Random Forest, Gradient Boosting, and Logistic Regression implementations in the scikit-learn Python package^31^. The hyperparameter values explored during tuning are reported in Supplementary Tables 1-4.

To benchmark classifier performance, we also computed the AUPRC for two control predictors. The first control predictor assigns a random binary label (0 or 1) to each data point, while the second predicts the negative class for all data points.

## Results

We used Principal Component Analysis (PCA), as implemented in the scikit-learn Python package^31^, to perform dimensionality reduction and visualize TCR sequence embeddings. The Full dataset derived from VDJdb comprises 24,751 TCR-pMHC pairs. A subset comprising 13,366 TCRs bind KLGGALQAK-A*03:01 and constitute the KLG positive set, while 10,035 TCRs do not bind KLGGALQAK-A*03:01 and form the corresponding negative set. In addition, 947 TCRs bind KLGGALQAK-A*03:01 as well as at least one other pMHC. Visualization of the ESM-2 embeddings reveals substantial overlap between positive and negative TCRs despite the PCA revealing two clusters (Figure 1a), as well as between positive TCRs and healthy background TCRs spread across four clusters (Figure 1b).

**Figure 1.**
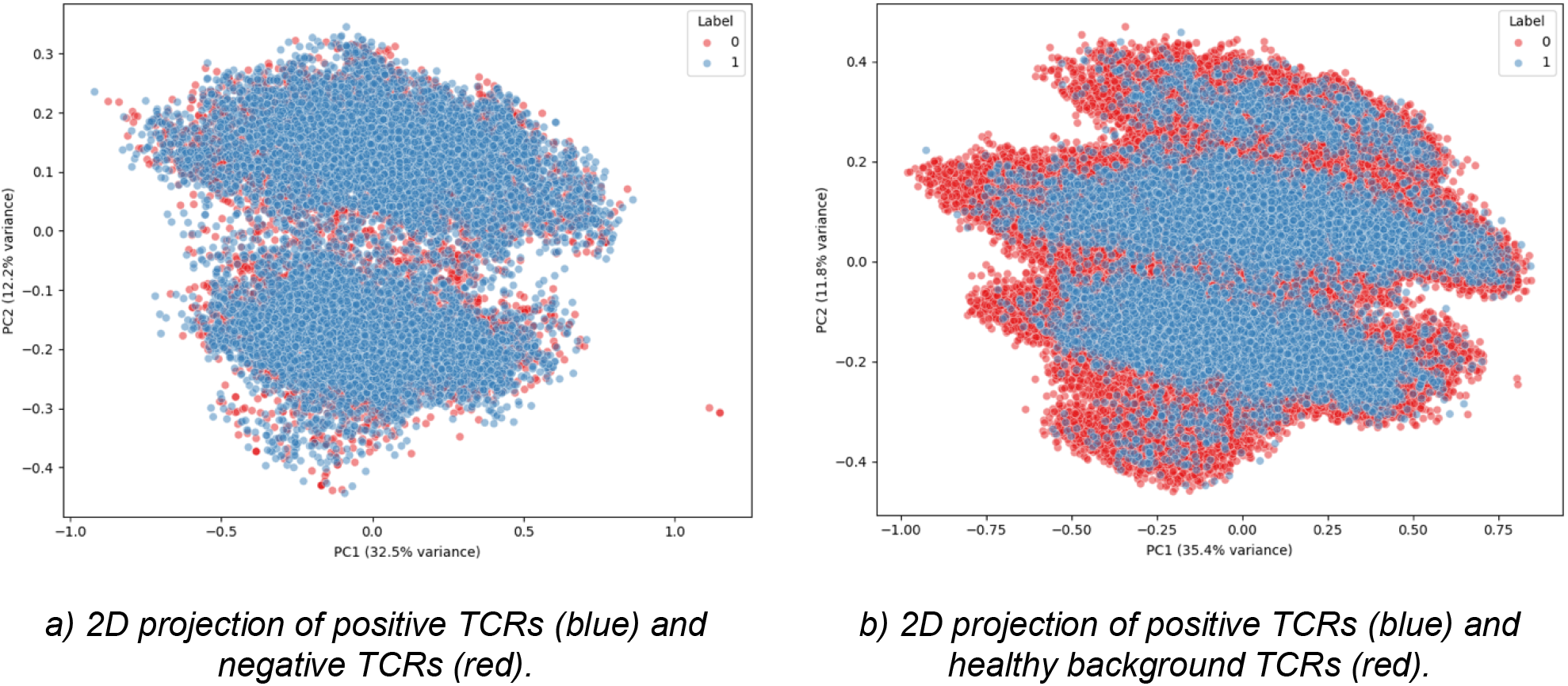
Two-dimensional visualization of the ESM-2 embeddings of TCRs binding KLGGALQAK-A*03:01 (positive TCRs) and corresponding negative TCRs. ESM-2 embeddings were projected into two dimensions using PCA.

Of the Full TCR set, 1,892 TCRs bind to GILGFVFTL-A*02:01, making our GIL positive set. After excluding 68 TCRs that bind GILGFVFTL-A*02:01 as well as one additional pMHC, the corresponding negative set comprises 22,486 TCRs. The overlap between positive and negative TCRs is shown in Figures 2a and 2b.

**Figure 2.**
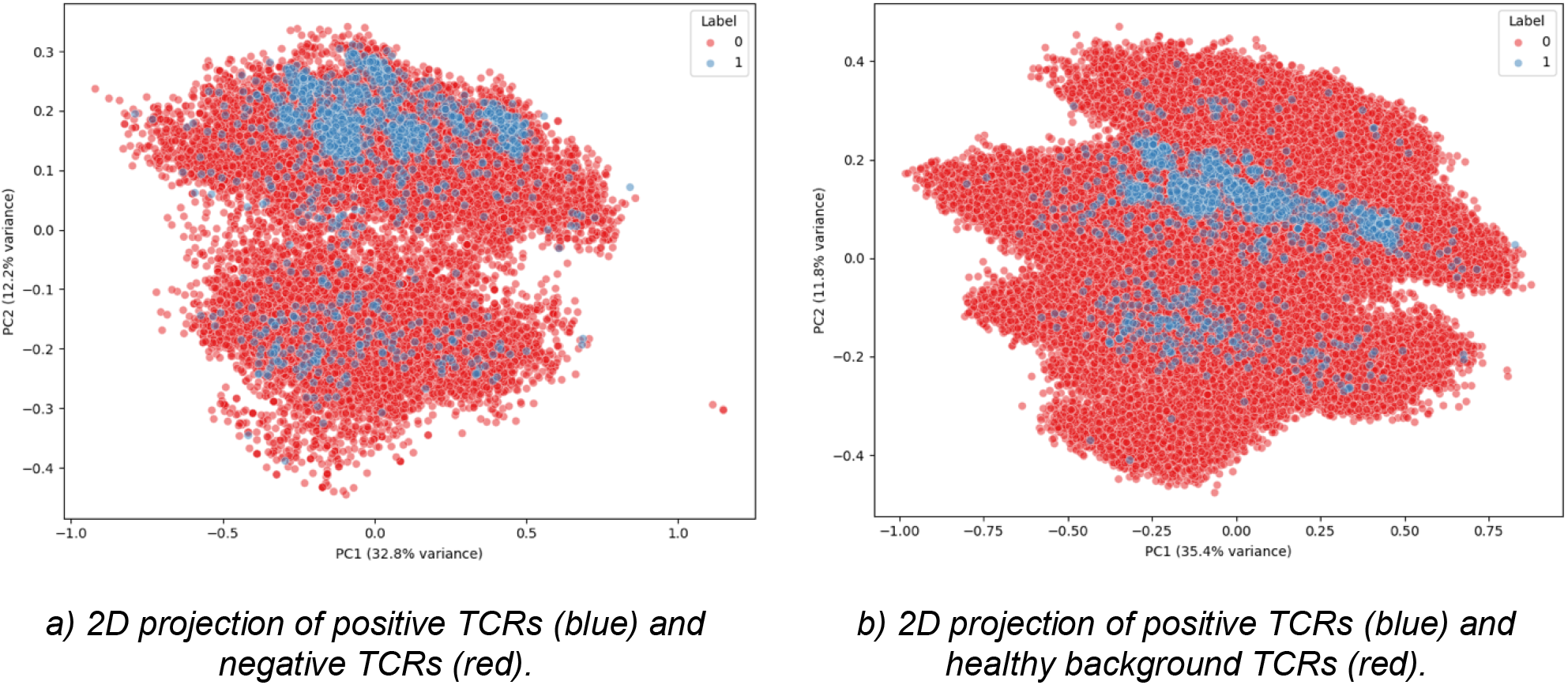
Two-dimensional visualization of ESM-2 embeddings for TCRs binding GILGFVFTL-A*02:01 (positive TCRs) and corresponding negative TCRs. ESM-2 embeddings were projected into two dimensions using PCA.

We also generated sequence embeddings using only the CDR3 regions of the TCR sequences using TCR-BERT and used PCA to perform dimensionality reduction and visualize these embeddings. For both KLGGALQAK-A*03:01 and GILGFVFTL-A*02:01, we can see overlap between the embeddings of the positive TCRs and the negative TCRs (Figures 3a and 4a) as well as the positive and healthy background TCRs (Figures 3b and 4b).

**Figure 3.**
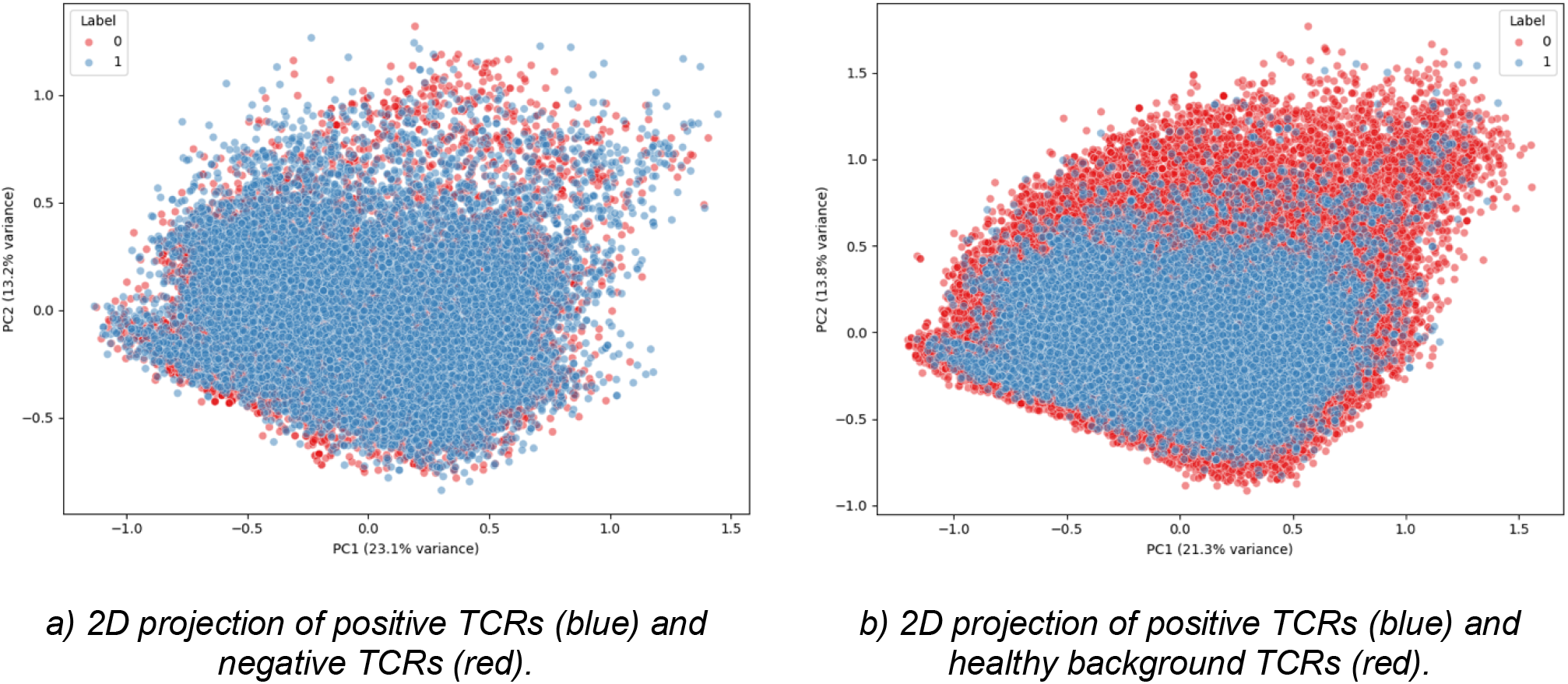
Two-dimensional PCA visualization of TCR-BERT embeddings of the CDR3 regions for TCRs binding KLGGALQAK-A*03:01 (positive TCRs) and corresponding negative TCRs.

**Figure 4.**
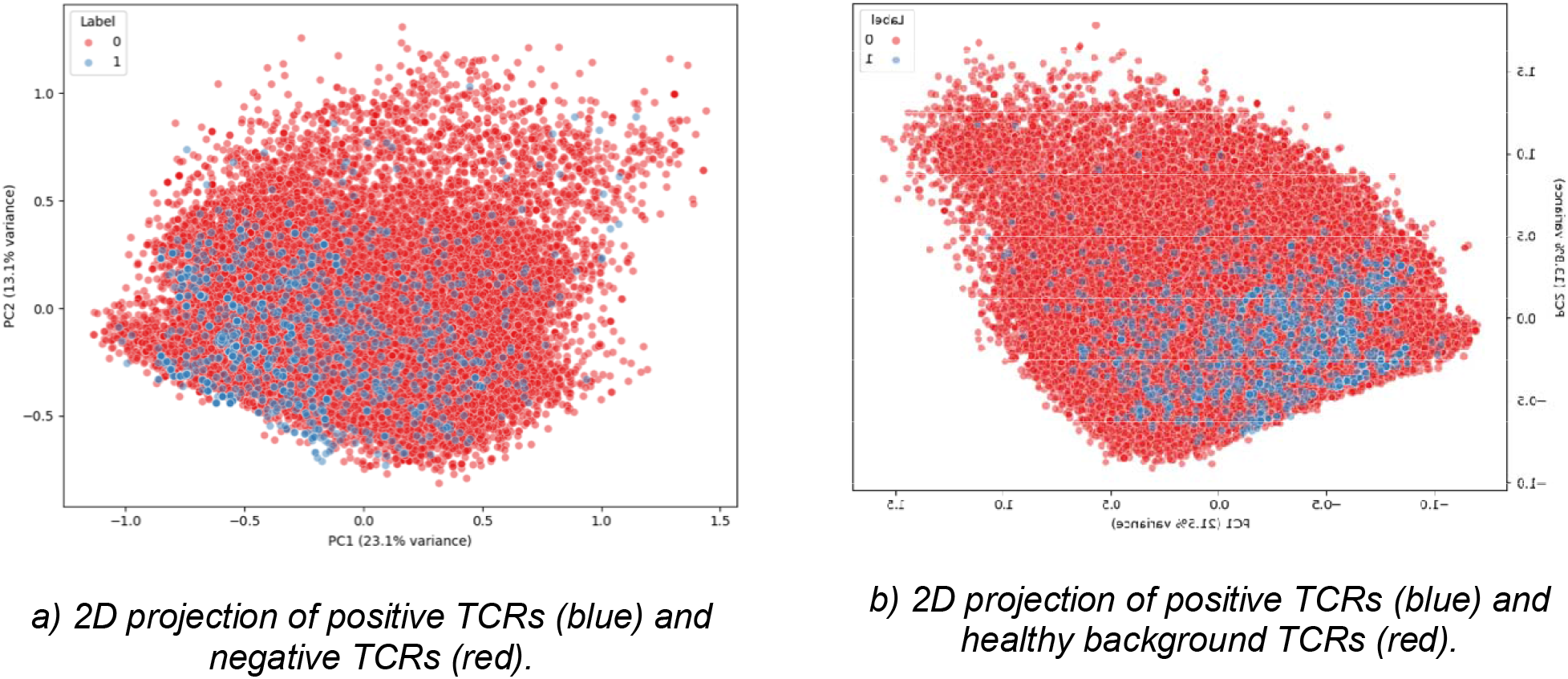
Two-dimensional PCA visualization of TCR-BERT embeddings of the CDR3 regions for TCRs binding GILGFVFTL-A*02:01 (positive TCRs) and corresponding negative TCRs.

We performed similar dimensionality reduction and visualization with t-SNE, as implemented in scikit-learn. The t-SNE visualizations of ESM-2 and TCR-BERT embeddings also show significant overlap in positive and negative TCRs and positive and healthy TCRs, as seen in Supplementary Figures 1-4.

Finally, we trained and evaluated Random Forest classifiers using each TCR sequence embeddings as input features. For KLGGALQAK-A*03:01, classifiers trained on most embeddings perform slightly worse than a baseline predictor that always predicts the negative class (Fig. 5a). When healthy background TCRs are used as negatives, performance further declines and falls below that of a random predictor (Fig. 5b).

**Figure 5.**
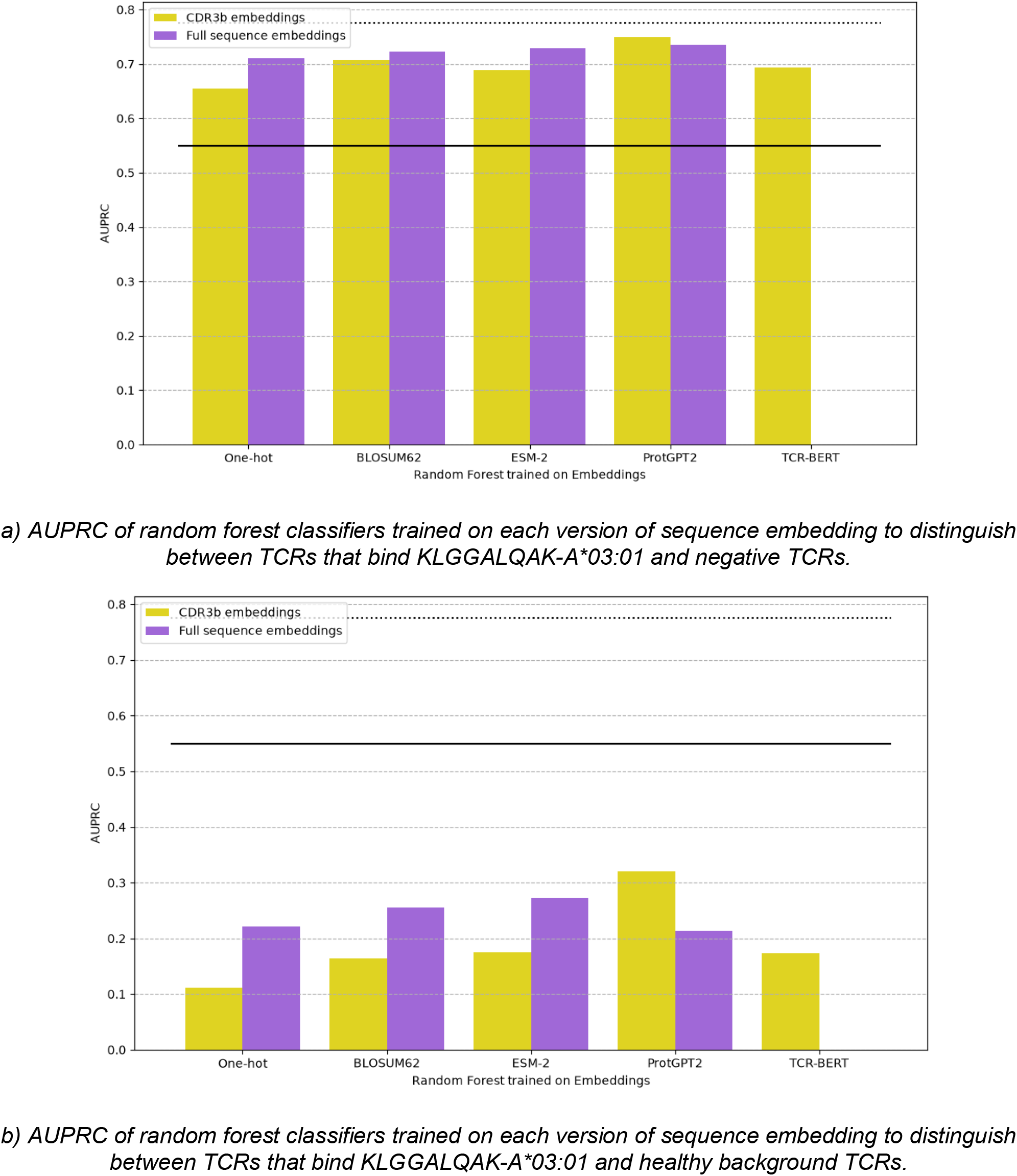
Area under the precision-recall curve (AUPRC) for classifiers trained to distinguish TCRs binding KLGGALQAK-A*03:01 from negative or healthy background TCRs. Solid black lines indicate the AUPRC of a random predictor, and dotted black lines indicate the AUPRC of a predictor that always predicts the negative class. Random Forest classifiers were trained on the sequence embeddings.

In contrast, for GILGFVFTL-A*02:01, all random forest classifiers outperform both the random and negative control predictors. The same trend is observed when healthy background TCRs are used as the negative set (Fig. 6).

**Figure 6.**
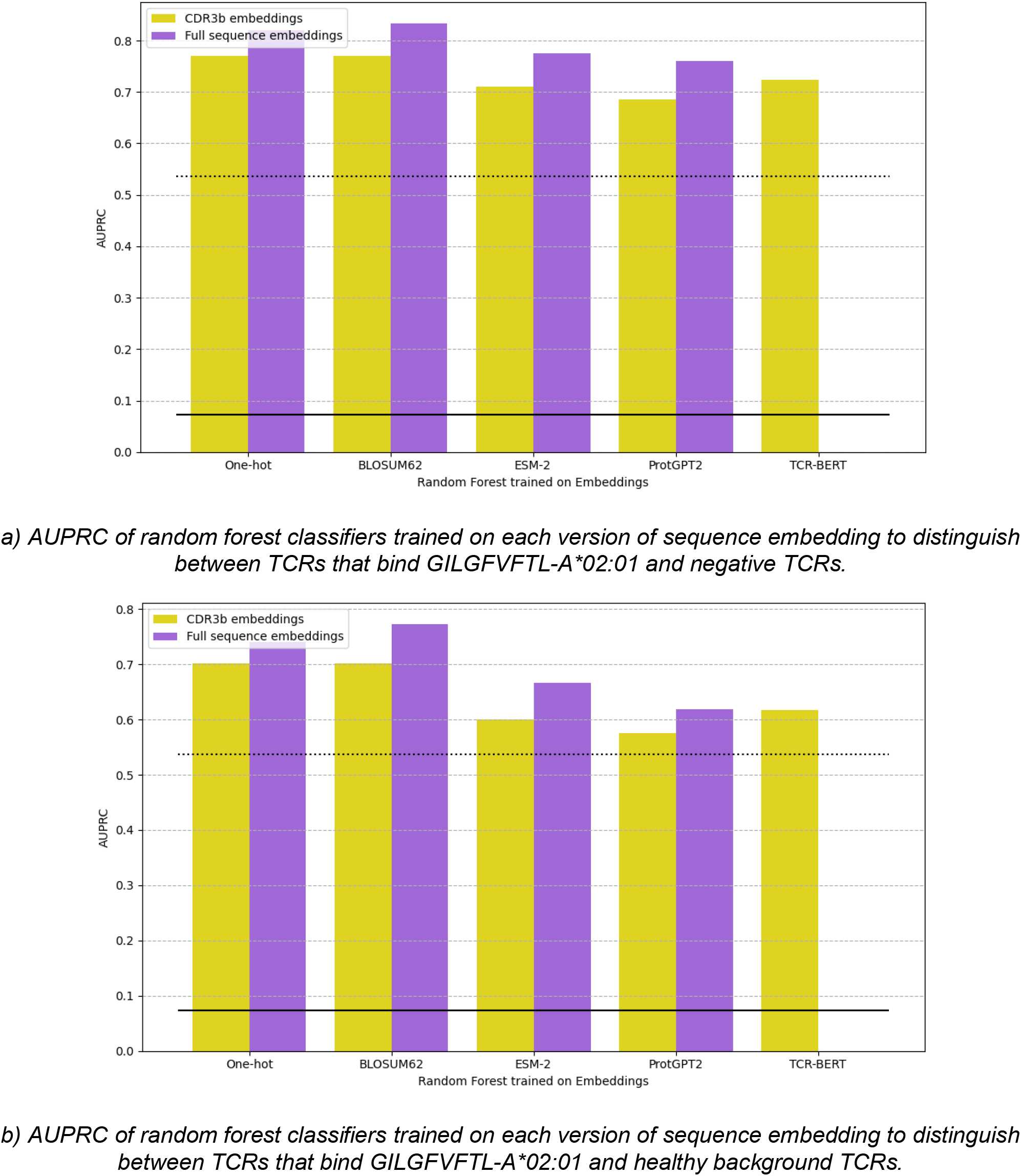
Area under the precision-recall curve (AUPRC) for classifiers trained to distinguish TCRs binding GILGFVFTL-A*02:01 from negative or healthy background TCRs. Solid black lines indicate the AUPRC for a random predictor, and dotted black lines indicate the AUPRC of a predictor that always predicts the negative class. Random Forest classifiers were trained on the sequence embeddings.

We trained additional classifiers (Gradient Boosting, MLP, and Logistic Regression) on the same task to assess how classifier choice affects performance. As shown in Supplementary Figures 5–7, performance on the KLG set was broadly consistent across classifiers, except for notably high performance by Logistic Regression and MLP models trained on full-sequence ESM-2 embeddings with healthy background TCRs as the negative set. Because this strong performance occurred only under this specific condition, it appears to be an outlier relative to other parameter settings. On the GIL set, Gradient Boosting performed worse than the baseline predictors when trained on ESM-2, ProtGPT2, and TCR-BERT embeddings (Supplementary Figure 8). MLP and Logistic Regression followed trends similar to Random Forest, as shown in Supplementary Figures 9 and 10.

## Discussion

Visualization of ESM-2 embeddings reveals substantial overlap between positive and negative data points, but, depending on the peptide, a random forest classifier trained on these embeddings can differentiate between positive and negative points with varying success. For GILGFVFTL-A*02:01, the random forest classifiers outperform both random and the constant baselines, whereas for KLGGALQAK-A*03:01, performance is comparable to or worse than the control classifiers. This difference may reflect greater similarity between TCRs that bind GILGFVFTL-A*02:01 compared with those that bind KLGGALQAK-A*03:01.

Several recent TCR specificity prediction methods have adopted ESM-2 embeddings as input features. In our study, we use Random Forest classifiers trained on simple sequence encodings (one-hot encoding and BLOSUM62) with classifiers trained on general protein language model embeddings (ESM-2 and ProtGPT2). Classifiers trained on simple embeddings perform comparable to, or slightly better than, those trained on general protein language model embeddings, suggesting that the latter do not capture important features more effectively than the simple one-hot or BLOSUM encodings.

These findings are consistent with recent benchmarking studies showing that most top-performing TCR specificity prediction models rely on hand-crafted features or simple encodings, and that general protein language model embeddings do not consistently improve predictive performance^13^. In addition, the classifier performance does not improve when trained on TCR-BERT embeddings, even though TCR-BERT is a TCR-specific model that only uses the CDR3*β* regions, which is known to play a central role in antigen recognition and is therefore the focus of many TCR specificity prediction methods. Similarly, restricting other embedding methods to the CDR3*β* region does not improve performance relative to using the full TCR sequence.

We next examined how the choice of negative data affects classifier performance. For all Random Forest classifiers trained on general sequence embeddings (ESM-2, ProtGPT2, one-hot, and BLOSUM62) performance decreases when distinguishing between positive TCRs and the healthy background set compared to the negative set from the Full dataset. This reduced performance likely reflects greater similarity between positive TCRs and healthy background TCRs, as well as increased class imbalance introduced by the larger background set.

The decreased performance is more pronounced for the KLG positive set than for the GIL positive set, suggesting that the KLG-specific TCRs are similar to the healthy background TCRs than the GIL-specific TCRs. In addition, the larger size of the KLG positive set compared to the GIL positive set, likely contributes to greater TCR diversity, which may further limit the ability of the Random Forest classifier to distinguish positive from negative TCRs.

Overall, our results highlight a limitation of sequence-based TCR embeddings derived from general protein language models in their ability to distinguish between positive and negative TCRs. ESM-2 and ProtGPT2 are trained on a large corpus of protein sequences, which allows them to learn the constraints that guide protein evolution.

Thus, these general protein language models are able to accurately predict protein function and even generate new evolutionarily consistent sequences^21,22^. However, the CDR regions of TCRs are not evolutionarily constrained. A study addressing the hypervariability of the CDR regions of antibodies proposed a model that refines foundation protein language models with a framework that focuses on the hypervariable regions and found improved performance in antibody prediction tasks^32^. Antibodies undergo a similar V(D)J recombination as TCRs to produce the hypervariable CDRs, so a similar TCR-specific method may be designed to address the lack of evolutionary constraints on the CDR. A study on the use of PLMs on B cell receptor specificity data notes that PLMs tend to underperform on low-sequence diversity datasets, but perform well on high-sequence diversity datasets^33^.

This limitation may also be alleviated by the use of more advanced models that more explicitly represent TCR-pMHC binding mechanisms, such as structural models or methods that incorporate biophysical properties of the TCR and pMHC. Previous studies have explored the potential of structural modeling for TCR specificity prediction^16,34^.

Model performance may also be limited by the absence of experimentally validated negative examples in available data sets. In addition, TCR specificity assays are known to exhibit high false-negative rates, particularly for low-affinity TCRs^35^. This further complicates the issue of false negatives in training data, especially when “negatives” are selected from the same study as the positives. While some studies have reported advantages to using background TCR repertoires from healthy donors^15^, others have noted potential confounding effects arising from differences in donor populations and experimental methods^36^. Another study notes that model performance improves the more similar the positive and negative training datasets are to each other^37^.

Public databases also suffer from limited peptide diversity. As noted previously, approximately 100 epitopes account for nearly 70% of TCR-epitope pairs in public repositories^38^. The Full dataset from VDJdb used in this work contains 800 unique peptide-MHC pairs. The average number of TCRs per peptide-MHC is 31.04 but the median is 2, because 394 of the peptide-MHCs only have 1 associated positive TCR. This strong imbalance has been shown to limit model generalization, with even high performing models failing to predict specificity for unseen or underrepresented peptides^12–14,39^. Consequently, the lack of peptide diversity presents an additional challenge for training generalizable TCR specificity models, as models may overfit to a small number of overrepresented epitopes.

In this study, we identify fundamental issues preventing advances in the field of TCR specificity prediction as 1) the inability of general protein language models and sequence embedding techniques to distinguish between positive and negative TCRs and 2) the lack of experimentally validated negative examples in public TCR specificity data. Future work should prioritize the generation of experimentally validated negative examples, as well as computational methods that explicitly account for TCR-pMHC binding mechanisms. In addition, focusing on epitope-specific (local) models rather than global models trained across all epitopes may improve performance for underrepresented peptides. Finally, further investigation of structure-based embeddings may offer additional gains in TCR specificity prediction.

## Acknowledgements

We would like to thank Rob Stanton, Chirag Krishna, Kai-chi Huang, Xinli Hu, and Mojtaba Haghighatlari for their useful comments and guidance.

## References

1. Janeway, C. A., Travers, P., Walport, M. & Shlomchik, M. J. The major histocompatibility complex and its functions. in Immunobiology: The Immune System in Health and Disease. 5th edition (Garland Science, 2001).

2. Ura, T. et al. Current Vaccine Platforms in Enhancing T-Cell Response. Vaccines 10, 1367 (2022).

3. Pollard, A. J. & Bijker, E. M. A guide to vaccinology: from basic principles to new developments. Nat. Rev. Immunol. 21, 83–100 (2021).

4. Waldman, A. D., Fritz, J. M. & Lenardo, M. J. A guide to cancer immunotherapy: from T cell basic science to clinical practice. Nat. Rev. Immunol. 20, 651–668 (2020).

5. Seiringer, P., Garzorz-Stark, N. & Eyerich, K. T-Cell_Mediated Autoimmunity: Mechanisms and Future Directions. J. Invest. Dermatol. 142, 804–810 (2022).

6. Arstila, T. P. et al. A Direct Estimate of the Human *αβ* T Cell Receptor Diversity.Science 286, 958–961 (1999).

7. Nikolich-Žugich, J., Slifka, M. K. & Messaoudi, I. The many important facets of T-cell repertoire diversity. Nat. Rev. Immunol. 4, 123–132 (2004).

8. Goncharov, M. et al. VDJdb in the pandemic era: a compendium of T cell receptors specific for SARS-CoV-2. Nat. Methods 19, 1017–1019 (2022).

9. Tickotsky, N., Sagiv, T., Prilusky, J., Shifrut, E. & Friedman, N. McPAS-TCR: a manually curated catalogue of pathology-associated T cell receptor sequences. Bioinformatics 33, 2924–2929 (2017).

10. Nolan, S. et al. A large-scale database of T-cell receptor beta sequences and binding associations from natural and synthetic exposure to SARS-CoV-2. Front. Immunol. 16, 1488851 (2025).

11. Vita, R. et al. The Immune Epitope Database (IEDB): 2018 update. Nucleic Acids Res. 47, D339–D343 (2019).

12. Nielsen, M. et al. Lessons learned from the IMMREP23 TCR-epitope prediction challenge. ImmunoInformatics 16, 100045 (2024).

13. Deng, L. et al. Performance comparison of TCR-pMHC prediction tools reveals a strong data dependency. Front. Immunol. 14, 1128326 (2023).

14. Montemurro, A. et al. NetTCR-2.0 enables accurate prediction of TCR-peptide binding by using paired TCR*α* and *β* sequence data. Commun. Biol. 4, 1060 (2021).

15. Zhang, J., Ma, W. & Yao, H. Accurate TCR-pMHC interaction prediction using a BERT-based transfer learning method. Brief. Bioinform. 25, bbad436 (2023).

16. Slone, J. K. et al. STAG-LLM: Predicting TCR-pHLA binding with protein language models and computationally generated 3D structures. Comput. Struct. Biotechnol. J. 27, 3885–3896 (2025).

17. Luu, A., Leistico, J., Miller, T., Kim, S. & Song, J. Predicting TCR-Epitope Binding Specificity Using Deep Metric Learning and Multimodal Learning. Genes 12, 572 (2021).

18. Jokinen, E., Huuhtanen, J., Mustjoki, S., Heinonen, M. & Lähdesmäki, H. Predicting recognition between T cell receptors and epitopes with TCRGP. PLOS Comput. Biol. 17, e1008814 (2021).

19. Yang, M. et al. MIX-TPI: a flexible prediction framework for TCR–pMHC interactions based on multimodal representations. Bioinformatics 39, btad475 (2023).

20. Henikoff, S. & Henikoff, J. G. Amino acid substitution matrices from protein blocks. Proc. Natl. Acad. Sci. 89, 10915–10919 (1992).

21. Lin, Z. et al. Evolutionary-scale prediction of atomic-level protein structure with a language model.

22. Ferruz, N., Schmidt, S. & Höcker, B. ProtGPT2 is a deep unsupervised language model for protein design. Nat. Commun. 13, 4348 (2022).

23. Yadav, S., Vora, D. S., Sundar, D. & Dhanjal, J. K. TCR-ESM: Employing protein language embeddings to predict TCR-peptide-MHC binding. Comput. Struct. Biotechnol. J. 23, 165–173 (2024).

24. Breiman, L. Random Forests. Mach. Learn. 45, 5–32 (2001).

25. Weber, A., Born, J. & Rodriguez Martínez, M. TITAN: T-cell receptor specificity prediction with bimodal attention networks. Bioinformatics 37, i237–i244 (2021).

26. Springer, I., Tickotsky, N. & Louzoun, Y. Contribution of T Cell Receptor Alpha and Beta CDR3, MHC Typing, V and J Genes to Peptide Binding Prediction. Front. Immunol. 12, 664514 (2021).

27. Quast, N. P. et al. T-cell receptor structures and predictive models reveal comparable alpha and beta chain structural diversity despite differing genetic complexity. 2024.05.20.594940 Preprint at 10.1101/2024.05.20.594940 (2024).

28. Heather, J. M. et al. Stitchr: stitching coding TCR nucleotide sequences from V/J/CDR3 information. Nucleic Acids Res. 50, e68 (2022).

29. Suzek, B. E., Huang, H., McGarvey, P., Mazumder, R. & Wu, C. H. UniRef: comprehensive and non-redundant UniProt reference clusters. Bioinformatics 23, 1282–1288 (2007).

30. Wu, K. et al. TCR-BERT: learning the grammar of T-cell receptors for flexible antigen-xbinding analyses. Preprint at 10.1101/2021.11.18.469186 (2021).

31. Pedregosa, F. et al. Scikit-learn: Machine Learning in Python. Mach. Learn.PYTHON.

32. Singh, R. et al. Learning the language of antibody hypervariability. Proc. Natl. Acad. Sci. 122, e2418918121 (2025).

33. Frolenkova, M. et al. Quantitative mapping of antigen specificity in adaptive immune repertoire embedding spaces. Preprint at 10.64898/2025.12.09.692930 (2025).

34. Bradley, P. Structure-based prediction of T cell receptor:peptide-MHC interactions. eLife 12, e82813 (2023).

35. Rius, C. et al. Peptide–MHC Class I Tetramers Can Fail To Detect Relevant Functional T Cell Clonotypes and Underestimate Antigen-Reactive T Cell Populations. J. Immunol. 200, 2263–2279 (2018).

36. Weber, A., Pélissier, A. & Rodríguez Martínez, M. T-cell receptor binding prediction: A machine learning revolution. ImmunoInformatics 15, 100040 (2024).

37. Ursu, E. et al. Training data composition determines machine learning generalization and biological rule discovery. Nat. Mach. Intell. 7, 1206–1219 (2025).

38. Hudson, D., Fernandes, R. A., Basham, M., Ogg, G. & Koohy, H. Can we predict T cell specificity with digital biology and machine learning? Nat. Rev. Immunol. 23, 511–521 (2023).

39. Moris, P. et al. Current challenges for unseen-epitope TCR interaction prediction and a new perspective derived from image classification. Brief. Bioinform. 22, bbaa318 (2021).

