## Supplemental for "Limitations of general protein language models and public TCRpMHC data in training specificity prediction models"

**^1^**Machine Learning and Computational Sciences, Pfizer Worldwide Research Development and Medical, Cambridge, MA, USA

**Supplementary Tables 1-4: Hyperparameter tuning values.** Hyperparameter tuning for each classifier was performed using a grid search over these values.

**ST 1. Logistic Regression**

| **Hyperparameter** | **Values** |
| --- | --- |
| C | 0.001, 0.01, 0.1, 1, 10, 100 |
| solver | lbfgs, liblinear |
| class_weight | None, balanced |
| max_iter | 1000 |

**ST 2. Random Forest**

| **Hyperparameter** | **Values** |
| --- | --- |
| n_estimators | 50, 100, 200 |
| max_depth | 5, 10, 20, None |
| min_samples_split | 2, 5, 10 |
| class_weight | None, balanced |

**ST 3. Gradient Boosting**

| **Hyperparameter** | **Values** |
| --- | --- |
| n_estimators | 100 |
| learning_rate | 0.1, 0.2 |
| max_depth | 3, 5 |
| subsample | 0.8, 1.0 |

**ST 4. MLP**

| **Hyperparameter** | **Values** |
| --- | --- |
| hidden_dims | (128,), (256,), (128, 64), (256, 128, 64) |
| dropout_rate | 0.1, 0.3, 0.5 |
| learning_rate | 1e-4, 1e-3, 1e-2 |
| batch_size | 64, 128 |
| max_epochs | 120 |
| patience | 10 |
| weight_decay | 1e-5, 1e-4, 1e-3 |

| Supp Figs 1-4, t-SNE projections  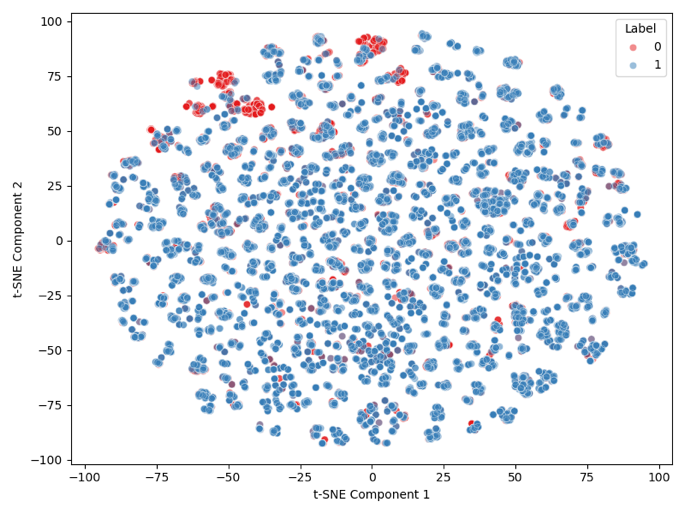  a) 2D projection of positive TCRs (blue) and negative TCRs (red). | 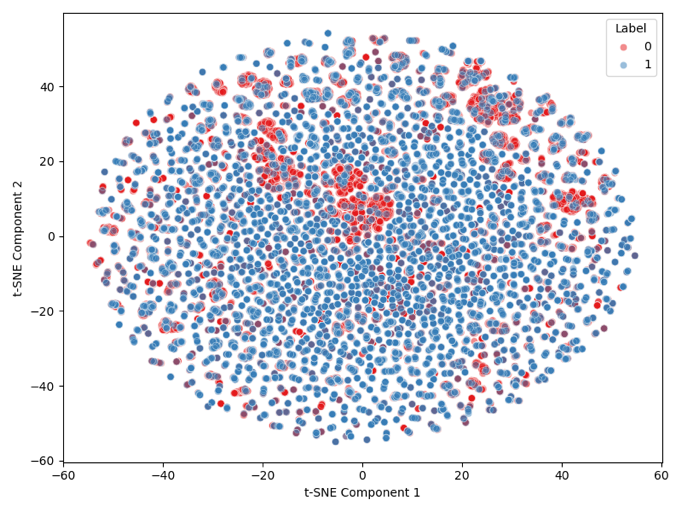  b) 2D projection of positive TCRs (blue) and healthy background TCRs (red). |
| --- | --- |

SF 1. Two-dimensional visualization of the ESM-2 embeddings of TCRs binding KLGGALQAK-A*03:01 (positive TCRs) and corresponding negative TCRs. ESM-2 embeddings were projected into two dimensions using t-SNE.

| 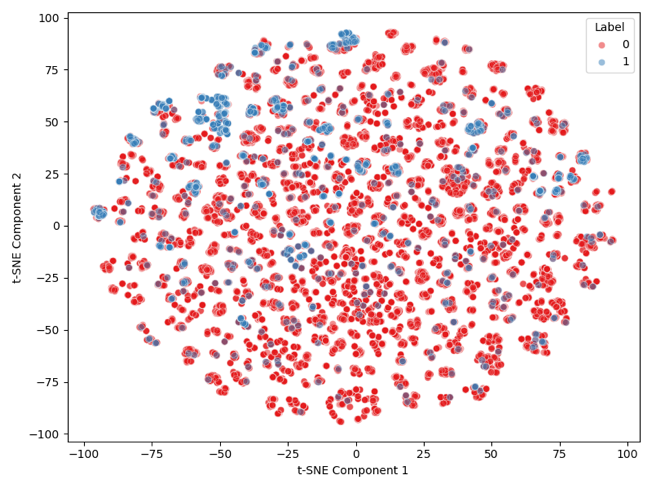  a) 2D projection of positive TCRs (blue) and negative TCRs (red). | 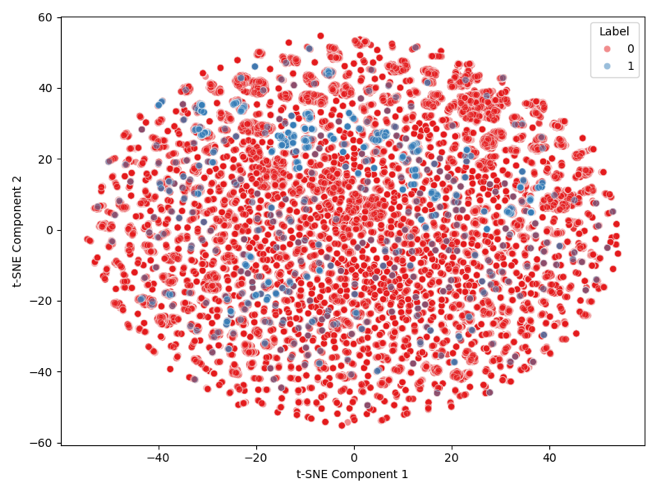  b) 2D projection of positive TCRs (blue) and healthy background TCRs (red). |
| --- | --- |

SF 2. Two-dimensional t-SNE visualization of ESM-2 embeddings for TCRs binding GILGFVFTL-A*02:01 (positive TCRs) and corresponding negative TCRs. ESM-2 embeddings were projected into two dimensions using t-SNE.

| 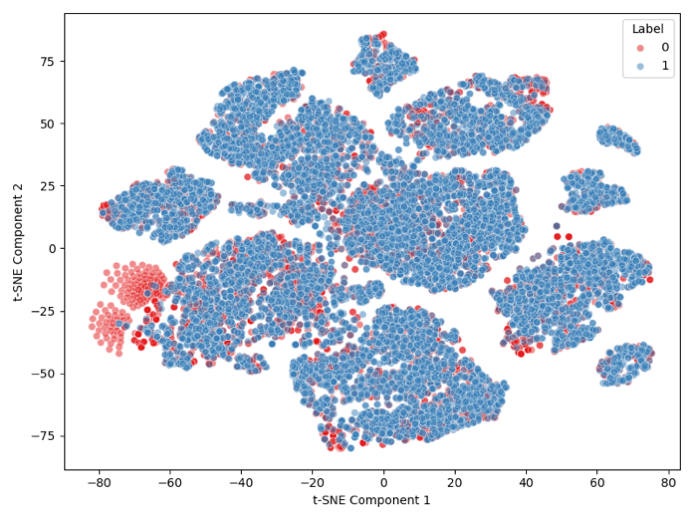  a) 2D projection of positive TCRs (blue) and negative TCRs (red). | 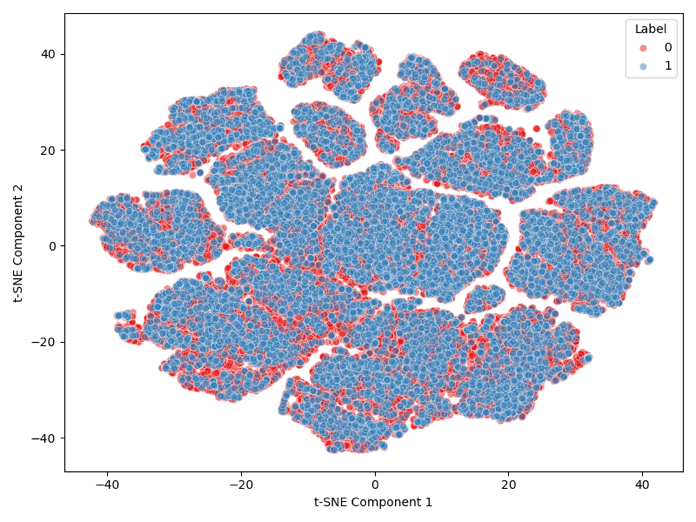  b) 2D projection of positive TCRs (blue) and healthy background TCRs (red). |
| --- | --- |

SF 3. Two-dimensional visualization of the ESM-2 embeddings of TCRs binding KLGGALQAK-A*03:01 (positive TCRs) and corresponding negative TCRs. ESM-2 embeddings were projected into two dimensions using t-SNE.

| 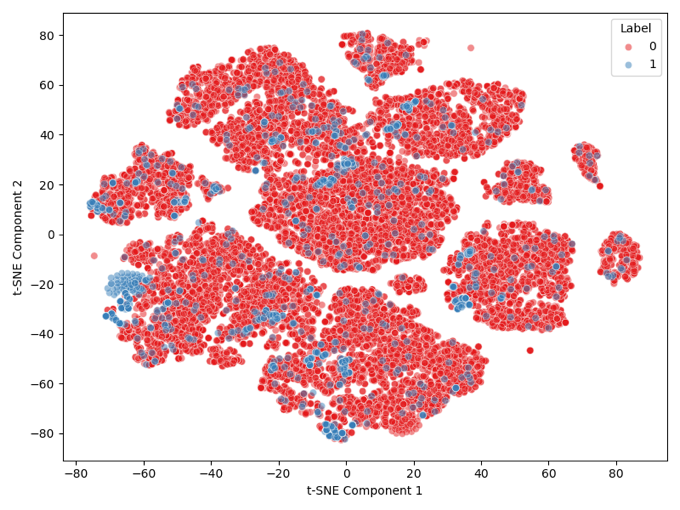  a) 2D projection of positive TCRs (blue) and negative TCRs (red). | 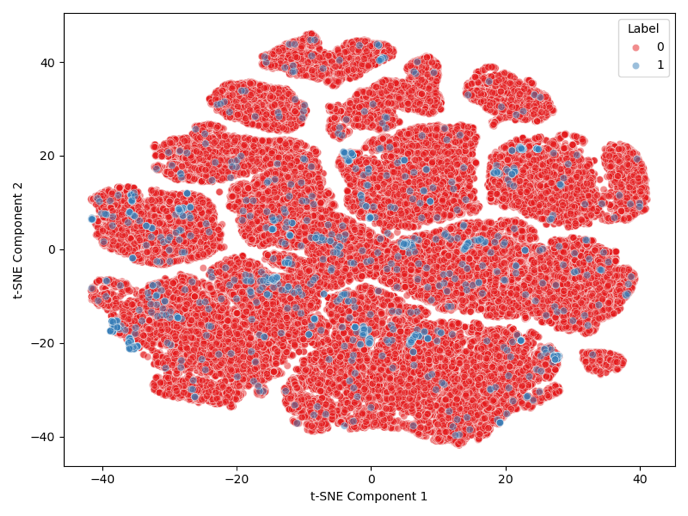  b) 2D projection of positive TCRs (blue) and healthy background TCRs (red). |
| --- | --- |

SF 4. Two-dimensional t-SNE visualization of ESM-2 embeddings for TCRs binding GILGFVFTL-A*02:01 (positive TCRs) and corresponding negative TCRs. ESM-2 embeddings were projected into two dimensions using t-SNE.

Supp Figures 5-7: Classifier performance on KLG set

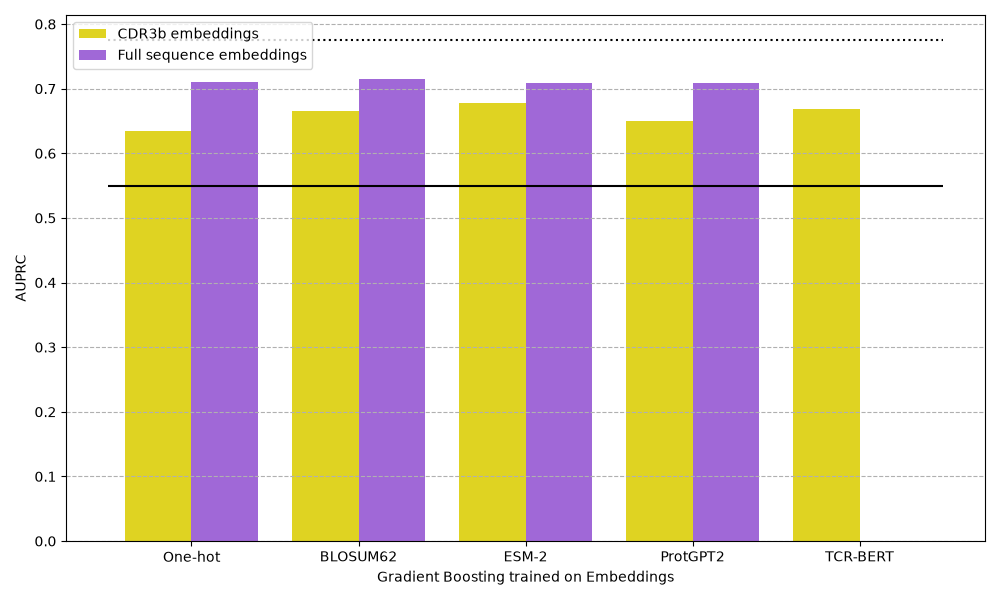

a) AUPRC of gradient boosting classifiers trained on each version of sequence embedding to distinguish between TCRs that bind KLGGALQAK-A*03:01 and negative TCRs.

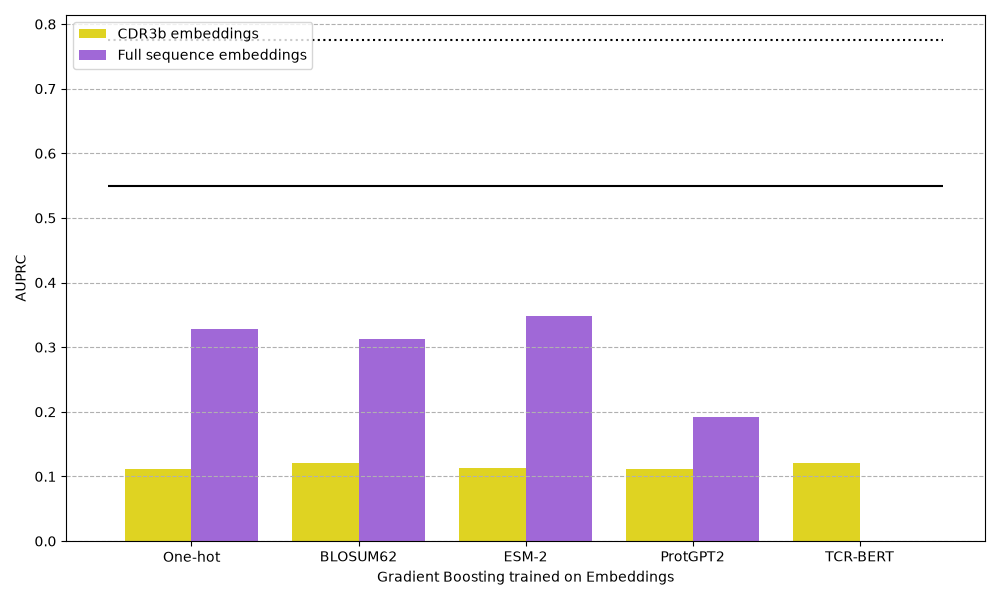

b) AUPRC of gradient boosting classifiers trained on each version of sequence embedding to distinguish between TCRs that bind KLGGALQAK-A*03:01 and healthy background TCRs.

SF 5. Area under the precision-recall curve (AUPRC) for classifiers trained to distinguish TCRs binding KLGGALQAK-A*03:01 from negative or healthy background TCRs. Solid black lines indicate the AUPRC of a random predictor, and dotted black lines indicate the AUPRC of a predictor that always predicts the negative class. Gradiant Boosting classifiers were trained on the sequence embeddings.

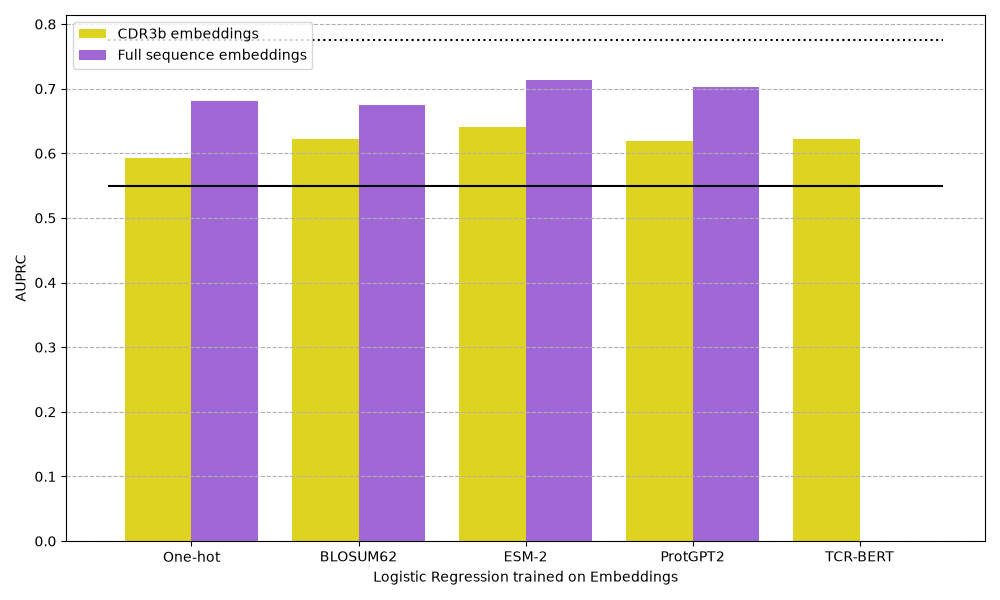

a) AUPRC of logistic regression classifiers trained on each version of sequence embedding to distinguish between TCRs that bind KLGGALQAK-A*03:01 and negative TCRs.

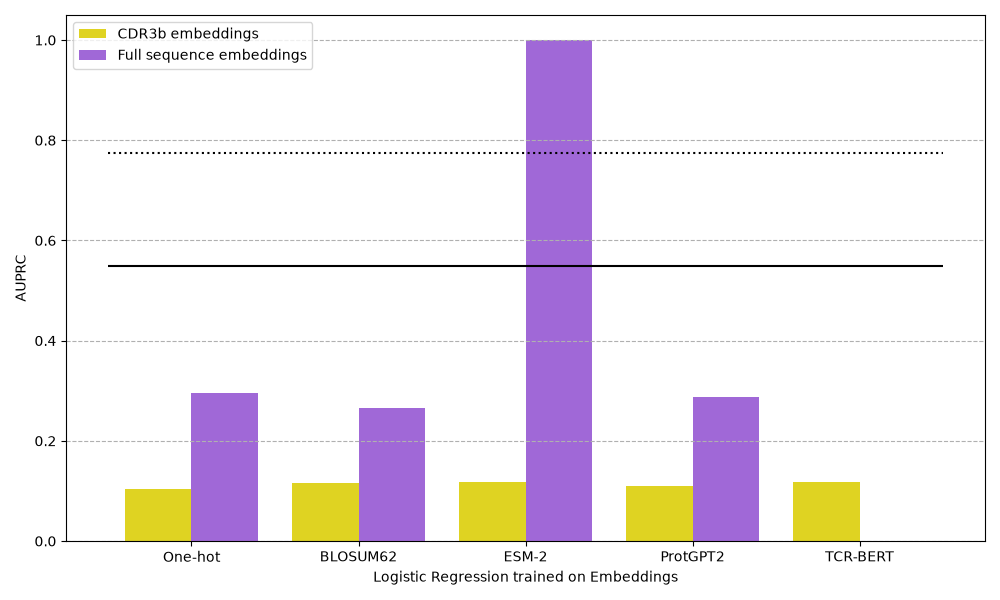

b) AUPRC of logistic regression classifiers trained on each version of sequence embedding to distinguish between TCRs that bind KLGGALQAK-A*03:01 and healthy background TCRs.

SF6. Area under the precision-recall curve (AUPRC) for classifiers trained to distinguish TCRs binding KLGGALQAK-A*03:01 from negative or healthy background TCRs. Solid black lines indicate the AUPRC of a random predictor, and dotted black lines indicate the AUPRC of a predictor that always predicts the negative class. Logistic Regression classifiers were trained on the sequence embeddings.

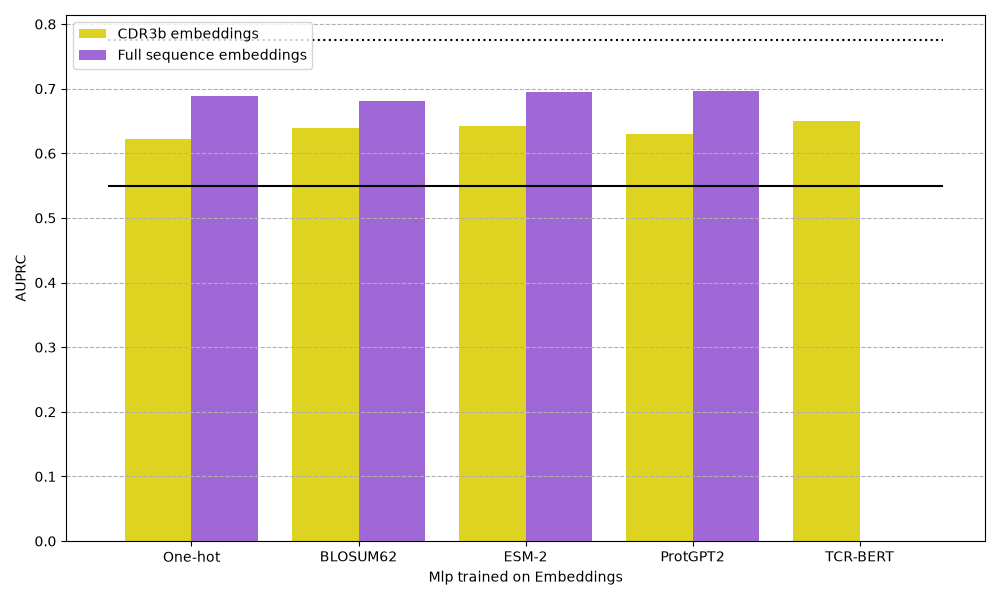

a) AUPRC of MLP classifiers trained on each version of sequence embedding to distinguish between TCRs that bind KLGGALQAK-A*03:01 and negative TCRs.

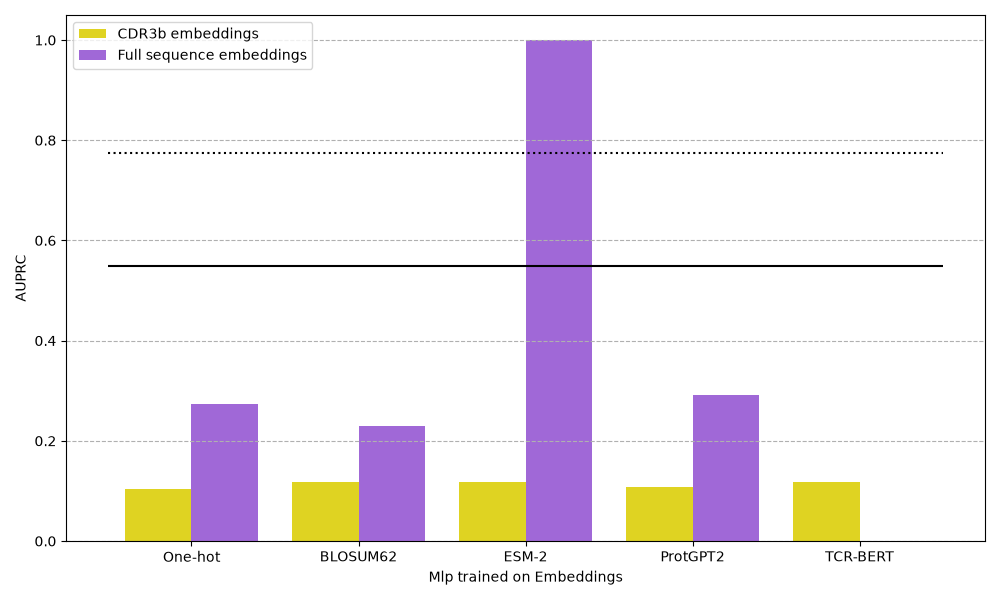

b) AUPRC of MLP classifiers trained on each version of sequence embedding to distinguish between TCRs that bind KLGGALQAK-A*03:01 and healthy background TCRs.

SF 7. Area under the precision-recall curve (AUPRC) for classifiers trained to distinguish TCRs binding KLGGALQAK-A*03:01 from negative or healthy background TCRs. Solid black lines indicate the AUPRC of a random predictor, and dotted black lines indicate the AUPRC of a predictor that always predicts the negative class.

Supp Figures 8-10: Classifier performance on GIL set

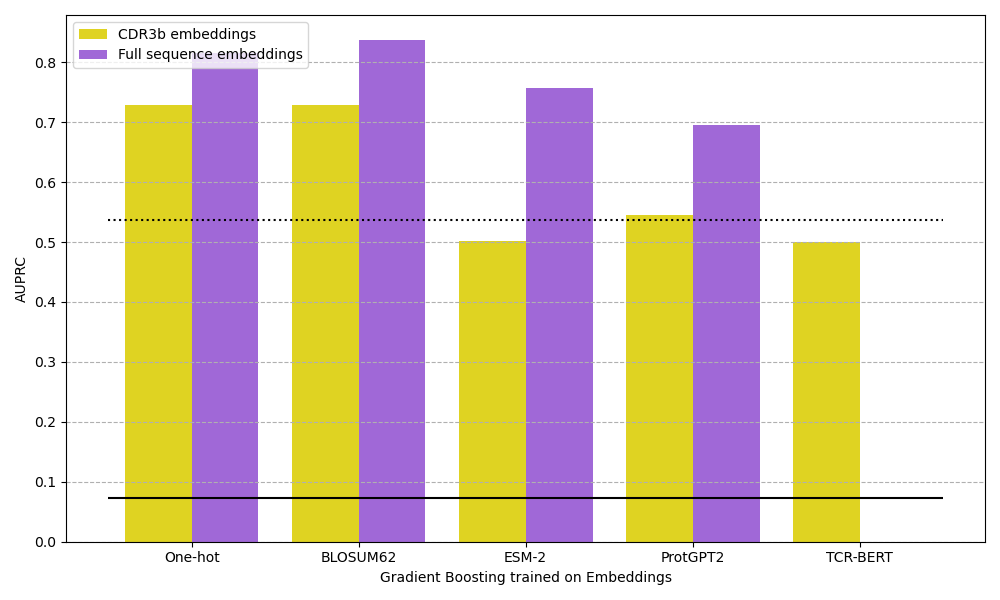

a) AUPRC of gradient boosting classifiers trained on each version of sequence embedding to distinguish between TCRs that bind GILGFVFTL-A*02:01and negative TCRs.

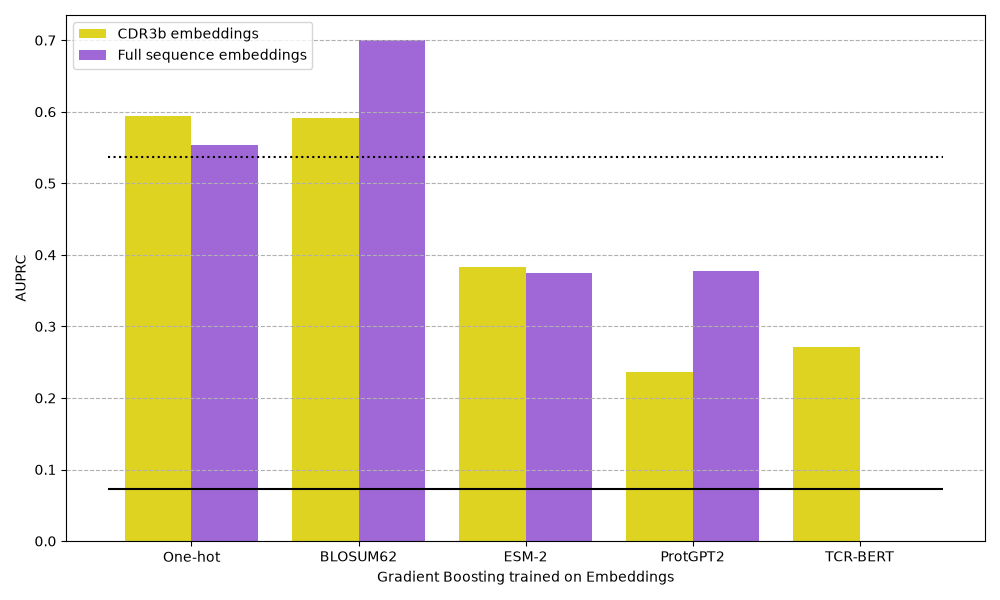

b) AUPRC of gradient boosting classifiers trained on each version of sequence embedding to distinguish between TCRs that bind GILGFVFTL-A*02:01and healthy background TCRs.

SF 8. Area under the precision-recall curve (AUPRC) for classifiers trained to distinguish TCRs binding GILGFVFTL-A*02:01 from negative or healthy background TCRs. Solid black lines indicate the AUPRC of a random predictor, and dotted black lines indicate the AUPRC of a predictor that always predicts the negative class. Gradiant Boosting classifiers were trained on the sequence embeddings.

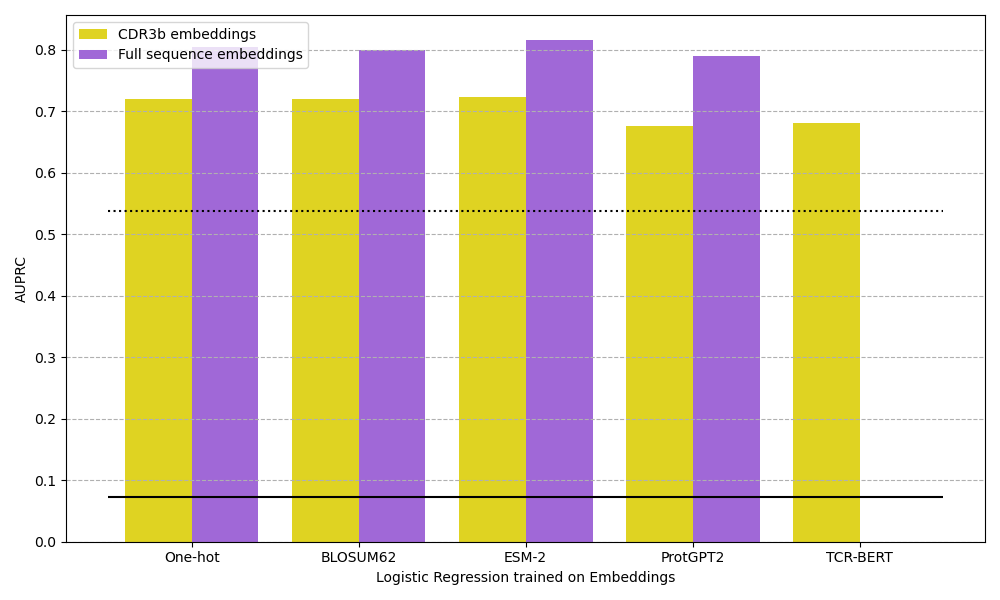

a) AUPRC of logistic regression classifiers trained on each version of sequence embedding to distinguish between TCRs that bind GILGFVFTL-A*02:01and negative TCRs.

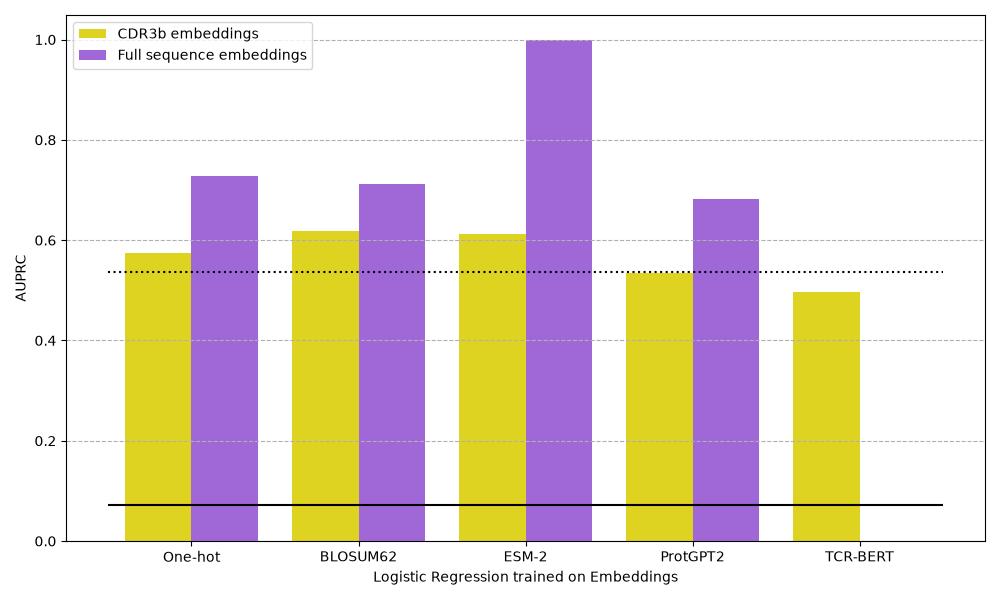

b) AUPRC of logistic regression classifiers trained on each version of sequence embedding to distinguish between TCRs that bind GILGFVFTL-A*02:01and healthy background TCRs.

SF 9. Area under the precision-recall curve (AUPRC) for classifiers trained to distinguish TCRs binding GILGFVFTL-A*02:01 from negative or healthy background TCRs. Solid black lines indicate the AUPRC of a random predictor, and dotted black lines indicate the AUPRC of a predictor that always predicts the negative class. Logistic Regression classifiers were trained on the sequence embeddings.

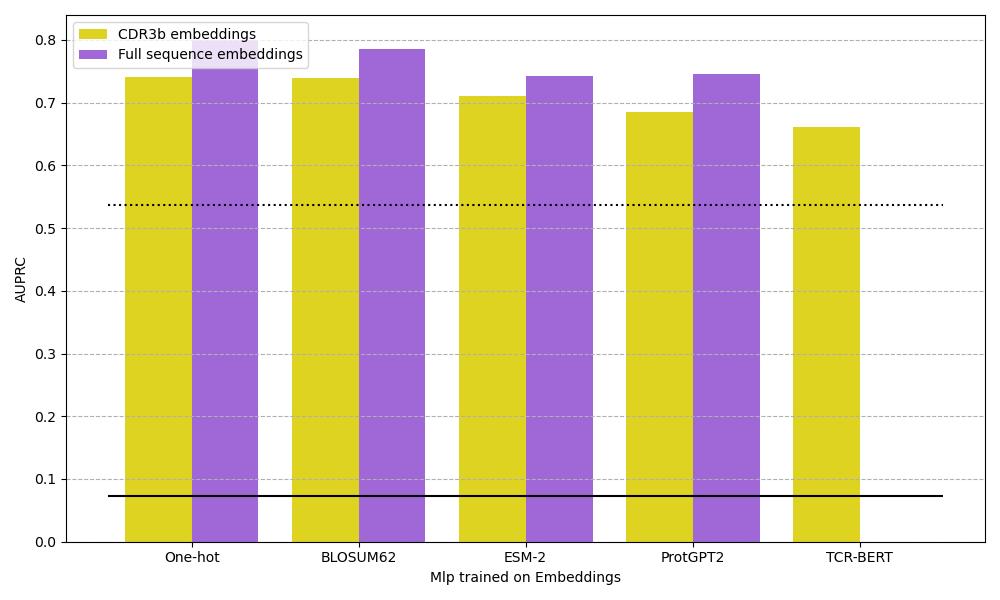

a) AUPRC of MLP classifiers trained on each version of sequence embedding to distinguish between TCRs that bind GILGFVFTL-A*02:01and negative TCRs.

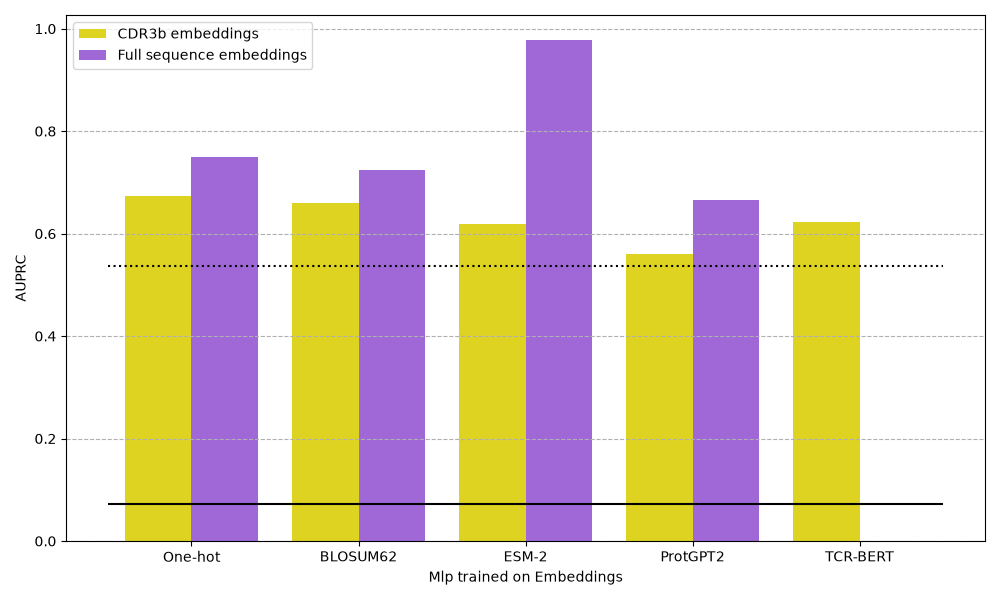

b) AUPRC of MLP classifiers trained on each version of sequence embedding to distinguish between TCRs that bind GILGFVFTL-A*02:01and healthy background TCRs.

SF 10. Area under the precision-recall curve (AUPRC) for classifiers trained to distinguish TCRs binding GILGFVFTL-A*02:01 and from negative or healthy background TCRs. Solid black lines indicate the AUPRC of a random predictor, and dotted black lines indicate the AUPRC of a predictor that always predicts the negative class.

   **Received:** , **Published:**
